# Flicker light stimulation induces thalamocortical hyperconnectivity with LGN and higher-order thalamic nuclei

**DOI:** 10.1101/2023.07.26.550646

**Authors:** Ioanna A. Amaya, Marianna E. Schmidt, Marie T. Bartossek, Johanna Kemmerer, Evgeniya Kirilina, Till Nierhaus, Timo T. Schmidt

## Abstract

The thalamus is primarily known as a relay for sensory information; however, it also critically contributes to higher-order cortical processing and coordination. Thalamocortical hyperconnectivity is associated with hallucinatory phenomena that occur in various psychopathologies (e.g., psychosis, migraine aura) and altered states of consciousness (ASC, e.g., induced by psychedelic drugs). However, the exact functional contribution of thalamocortical hyperconnectivity in forming hallucinatory experiences is unclear. Flicker light stimulation (FLS) can be used as an experimental tool to induce transient visual hallucinatory phenomena in healthy participants. Here, we use FLS in combination with fMRI to test how FLS modulates thalamocortical connectivity between specific thalamic nuclei and visual areas. We show that FLS induces thalamocortical hyperconnectivity between LGN, early visual areas and proximal upstream areas of ventral and dorsal visual streams (e.g., hV4, VO1, V3a). Further, an exploratory analysis indicates specific higher-order thalamic nuclei, such as anterior and mediodorsal nuclei, to be strongly affected by FLS. Here, the connectivity changes to upstream cortical visual areas directly reflect a frequency-dependent increase in experienced visual phenomena. Together these findings contribute to the identification of specific thalamocortical interactions in the emergence of visual hallucinations.

**Highlights:**

- Flicker light stimulation (FLS) induces thalamocortical hyperconnectivity between the first-order thalamic LGN and early visual cortices, likely due to entrainment.
- Thalamocortical connectivity between LGN and upstream visual areas, but not V1, is associated with the intensity of visual hallucinations.
- Thalamocortical connectivity changes with higher-order thalamic nuclei, such as anterior and mediodorsal nuclei, show strongest modulation by flicker frequency, which corresponds to the intensity of visual hallucinations.

## Introduction

The functional role of the thalamus goes beyond a relay for sensory information to the cortex. Indeed, no more than 20% of thalamic volume are primary sensory nuclei (Hádinger et al., 2023; Rovó et al., 2012). With complex connectivity throughout the neocortex, the thalamus contributes to higher-order processing, cognition and is also thought to coordinate information availability across cortices (Halassa and Sherman, 2019; Sherman and Guillery, 2006). Correspondingly, thalamocortical hyperconnectivity has been related to various pathologies, such as psychosis (Avram et al., 2021; Ramsay, 2019), epilepsy (Chen et al., 2021; Kim et al., 2014) and migraine (Bolay, 2020; Martinelli et al., 2021; Tu et al., 2019), and during diverse altered states of consciousness (ASC; e.g., induced by psychoactive drugs (Carhart-Harris et al., 2016; Müller et al., 2017; Preller et al., 2019)), all of which involve hallucinatory experiences (consider also (Hirschfeld et al., 2023; Hirschfeld and Schmidt, 2021; Prugger et al., 2022; Schmidt and Majić, 2017)). However, the exact functional contributions of thalamocortical hyperconnectivity to the emergence of hallucinatory phenomena is unclear. Previous reports are limited in the specificity of distinct thalamic nuclei contributions. Here, we utilize flicker light stimulation (FLS) in combination with fMRI to induce transient visual hallucinations in healthy participants and test for the differential modulation of functional connectivity between thalamic nuclei and visual areas.

The neural mechanisms of visual hallucinations are difficult to investigate empirically as their involvement in pathologies are spontaneous and co-exist with other neurophysiologic symptoms (Rogers et al., 2021). This makes it important to identify an experimental tool that can selectively induce visual hallucinatory phenomena in healthy participants. FLS applies stroboscopic light, primarily at alpha frequency (8-12 Hz), over closed eyes to elicit visual hallucinatory perception within seconds of stimulus onset. FLS-induced hallucinations include the perception of simple geometric patterns, motion and colours (Allefeld et al., 2011; Amaya et al., 2023; Bartossek et al., 2021; Montgomery et al., 2023), which hold close similarity to the content of visual hallucinations reported in migraine (Cowan, 2013; Panayiotopoulos, 1994; Richards, 1971; Schott, 2007; Wilkinson, 2004), epilepsy (Panayiotopoulos, 1994), psychedelic experiences (Bartossek et al., 2021; Klüver, 1966; Lawrence et al., 2022), and Charles Bonnet Syndrome (Ffytche, 2005; Jan and Castillo, 2012). FLS rhythmicity, frequency and brightness can be closely controlled in an experimental setting (Rogers et al., 2021), making it an optimal tool to investigate neural mechanisms of visual hallucinations.

By identifying which thalamic nuclei display altered connectivity with the cortex during visual hallucinations, the functional role of thalamocortical dysconnectivity can be indicated. The lateral geniculate nucleus (LGN) is the first-order thalamic nucleus for visual input and has bidirectional connections with V1. Here, feedforward thalamocortical projections relay visual information from the retina. Feedback corticogeniculate connections modulate activity of the LGN via inhibitory interneurons (Sherman and Guillery, 2006). These pathways determine LGN activity by streaming visual information (e.g., stimulus features (Andolina et al., 2007)) and integrating extra-visual modulations (e.g., attentional (Reinhold et al., 2023)) (see (Briggs, 2020) for review). The cortico-striato-thalamo-cortical (CSTC) model proposes that drug- and pathology-induced hallucinations arise from thalamocortical hyperconnectivity (Geyer and Vollenweider, 2008; Preller et al., 2019; Vollenweider and Geyer, 2001). With the perspective of the thalamus as a sensory gate, its contribution to hallucinations is mostly attributed to dysfunctional gating, leading to “sensory flooding” (Geyer and Vollenweider, 2008) and consequent cortical misinterpretation of sensory signals. In line with this suggestion, the LGN was found to have increased connectivity with the occipital cortex in patients with schizophrenia (Anticevic et al., 2014b) and during psychedelic experiences (Müller et al., 2018), which may reflect reduced thalamic gating capacities of the LGN to visual information passing to the cortex.

Recently, there has been more attention on the differential roles of first-order and higher-order thalamic nuclei in the generation of visual hallucinations (Vollenweider and Preller, 2020), which is facilitated by methodological advances allowing for parcellation of thalamic nuclei (Iglehart et al., 2020; Johansen-Berg et al., 2005). Higher-order nuclei do not receive input from sensory organs, instead, they orchestrate cortico-cortical communication and modulate activity of other thalamic nuclei (Sherman, 2016; Sherman and Guillery, 2006). With regards to visual processing, the inferior and lateral pulvinar are a group of higher-order nuclei with pronounced bidirectional anatomic connections to V1, V2 and V4 (Gattass et al., 2014; Shipp, 2003; Soares et al., 2001), contributing to visual processing and attention (Adams et al., 2000; Benevento and Rezak, 1976; Gattass et al., 2017; Guedj and Vuilleumier, 2020;

Kaas and Lyon, 2007; Saalmann et al., 2012). They are associated with the generation of hallucinatory phenomena as they show a reduction in volume, neuronal number, and neuronal size in individuals with schizophrenia (Byne et al., 2002; Danos et al., 2003) and dementia with Lewy Bodies (symptoms includes visual hallucination) (Erskine et al., 2017). In sum, the inferior and lateral pulvinar are candidate higher-order thalamic nuclei to contribute to the emergence of FLS-induced visual hallucinatory phenomena.

When aiming to identify the functional role of thalamocortical interactions in the emergence of visual hallucinations, it is relevant to test for differential contributions of visual stream areas with regards to their hierarchical organization. The visual cortex comprises of early visual cortices (EVC: V1-V3), which are typically defined by their retinotopic representation of the visual field (Engel et al., 1997; Sereno et al., 1995), and upstream visual areas, which show less pronounced retinotopy and are commonly described by their selective response to specific features of visual input, such as the activation preference for motion (hMT/V5; (Zeki et al., 1991)), shape (hV4/LO2; (Grill-Spector et al., 1998; Malach et al., 1995; Silson et al., 2013)), colour (hV4/VO1; (Persichetti et al., 2015)) and orientation (LO1; (Silson et al., 2013)). One previous EEG study indicated an increase in V4 activity during FLS (Ffytche, 2008), which may relate to the increased intensity of subjective experience of seeing shapes and colours. However, it is likely that altered processing in multiple visual areas relates to FLS-induced effects and the exact functional contributions of cortical areas along the visual hierarchy has not yet been reported.

In this study, we test whether FLS-induced visual hallucinations relate to altered thalamocortical connectivity, and which thalamic nuclei and visual areas are primarily modulated. We use constant light, 3 Hz FLS and 10 Hz FLS, expecting that 10 Hz FLS will induce stronger visual hallucinatory phenomena than 3 Hz FLS and constant light, as previously reported (See (Amaya et al., 2023; Bartossek et al., 2021)). We acquired resting state fMRI data and use the Automated Anatomical Labelling Atlas 3 (AAL3; (Rolls et al., 2020)) for thalamus parcellation and a volume-based maximum probability map (MPM) of visual topography (Wang et al., 2015) for parcellation of visual areas. We hypothesise that LGN will show hyperconnectivity with EVC for 3 Hz and 10 Hz FLS, as they receive excitatory signals from the retina and therefore synchronise to the periodic visual stimulus. For higher-order visual regions, as well as higher-order thalamic nuclei (e.g., inferior and lateral pulvinar), we expect to find parametric modulation of connectivity by the experimental conditions, such that constant light will induce hypoconnectivity, as found in previous work (Schmidt et al., 2020), and FLS will induce frequency-dependent increases in connectivity, whereby 10 Hz produces the strongest coupling. Thereby, changes in connectivity should resemble the intensity of subjective hallucination experience.

## Methods

### Participants

Twenty-four German speakers with no history of psychiatric or neurological disorders participated in the experiment (14 female; age range 20-41 years, mean (*M)* = 28 years standard deviation (*SD)* = 5.7 years). All participants were right-handed according to the Edinburgh Handedness Inventory (Oldfield, 1971) (mean laterality quotient = 77.1). Social media and student mailing lists were used for recruitment. Participants were informed about the study aims and background, such as possible risks of FLS, before giving written consent. The study was approved by the ethics committee of the Charité Universitätsmedizin Berlin (application number: EA4/143/18). All procedures were consistent with the guidelines included in the “Declaration of Helsinki – Ethical Principles for Medical Research Involving Human Subjects”.

### Flicker light stimulation

For presentation of the light stimulation, we used the light device Lucia N°03 (Light Attendance GmbH, Innsbruck, Austria), which has been developed to evoke hypnagogic visual impressions by intermittent light stimulation. It is equipped with one halogen lamp that is used for constant light stimulation and eight LEDs to apply FLS with high precision in timing and luminance via a programmable interface. Three light stimulation conditions were used: (1) Constant light stimulation at full intensity through a halogen lamp; (2) 3 Hz; and (3) 10 Hz FLS as 50% ON/ 50% OFF times with LED light at maximum intensity, as previously applied (Bartossek et al., 2021; Schwartzman et al., 2019). To apply light stimulation inside the fMRI scanner, the light device was mounted on an aluminium stand close to the end of the gantry at 150 ± 2cm from the participants’ eyes. To make the light stimulation comparable to a previous phenomenological study, where the lamp was positioned 50 cm from the participants’ eyes (Bartossek et al., 2021), two lenses were introduced into the MRI-mirror system to collect and focus the light [Figure 1A] to deliver approximately the same amount of light to the eyes. To protect the light device from overheating (as the in-built ventilation did not work in the magnetic field), a custom-made air cooling was used that comprised of an industrial vacuum cleaner positioned outside of the shielded MRI room to deliver cold air via extension hose to the light device.

**Figure 1.**
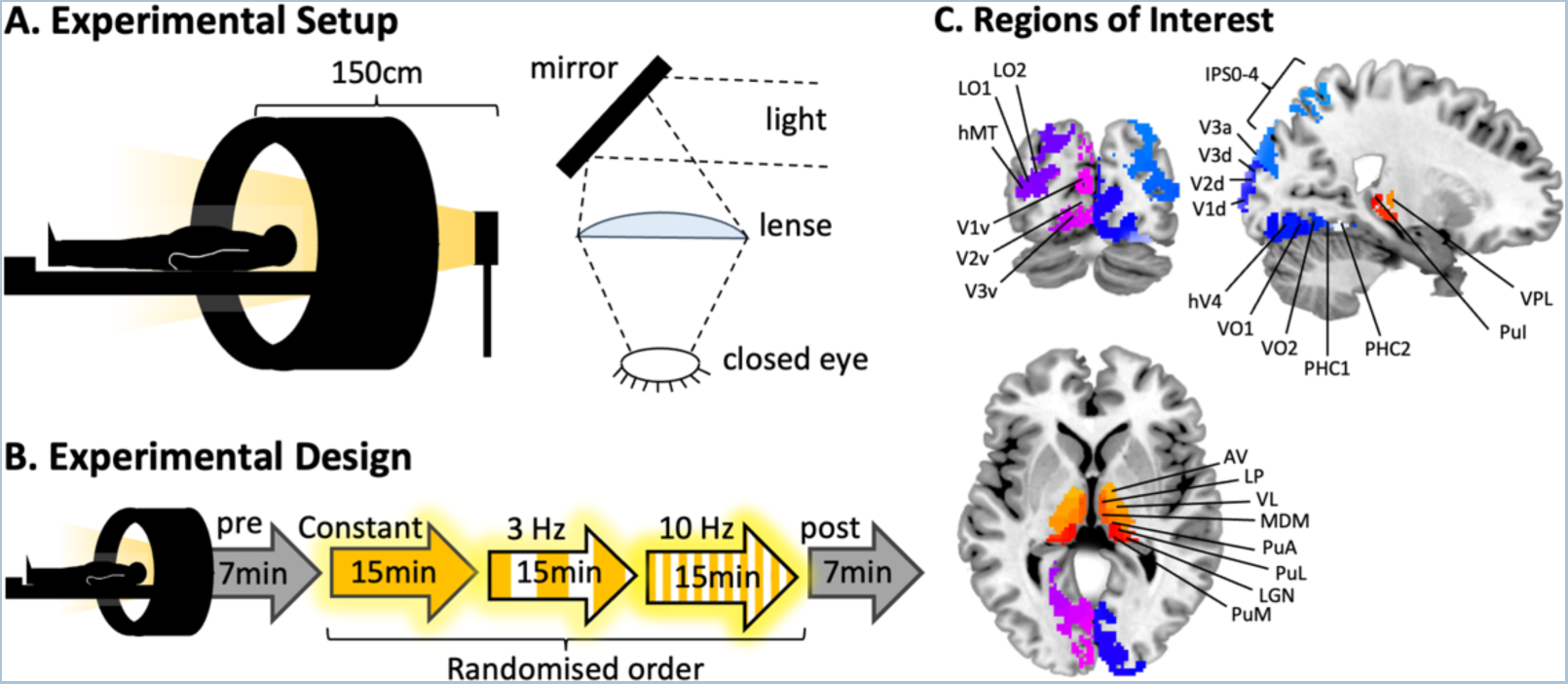
(A) Illustration of the setup inside the MRI scanner. FLS with the Lucia N°03 light device is optimized for stimulation from about 50 cm distance from the face. To obtain the same light intensity of stimulation inside of the scanner, the lamp was positioned at the end of the gantry at approximately 150 cm distance from the eyes and lenses were used to focus light onto the eyes to obtain the same amount of light as outside of the scanner. (B) The fMRI session comprised five closed-eye resting-state scans. The experimental conditions, constant light, 3 Hz and 10 Hz (15 minutes each) were presented in a randomised order, while the pre and post scans (7 minutes each) consisted of closed-eye rest for baseline measurements. (C) Regions of interest (ROIs) were extracted from AAL3 for thalamus parcellation (Rolls et al., 2020) and a volume-based MPM of visual topography (Wang et al., 2015). Thalamic ROIs, as labelled, are anteroventral (AV), lateroposterior (LP), ventrolateral (VL), mediodorsal medial (MDM), anterior pulvinar (PuA), lateral pulvinar (PuL), lateral geniculate nucleus (LGN), medial pulvinar (PuM), inferior pulvinar (PuI), and ventroposterolateral (VPL) nuclei. Additional thalamic ROIs not displayed are mediodorsal lateral (MDL), intralaminar (IL), ventroanterior (VA) and medial geniculate nuclei (MGN). Cortical ROIs of visual topography are split into the dorsal stream (V1d, V2d, V3d, V3a, V3b, LO1, LO2, hMT), ventral stream (V1v, V2v, V3v, hV4, VO1, VO2, PHC1, PHC2) and parietal stream (IPS0-4, SPL1, FEF). Cortical ROIs not displayed are V3b, SPL1 and FEF.

### Study Design and Procedure

To minimize risk of aversive effects of FLS, all participants underwent a preliminary semi-structured video-interview with a psychologist to identify any acute mental disorders, consumption of psychotropic medication and/or pregnancy. Thereafter, participants were screened for indications of photosensitive epilepsy based on electroencephalography (EEG) and were shortly presented FLS of each experimental condition to be familiarized with the procedures and setup. All measurements were conducted at the Center for Cognitive Neuroscience Berlin (CCNB) at the Freie Universität Berlin.

The scanning session comprised of five scans: pre and post scans, each lasting seven minutes and consisting of closed-eye rest in darkness, and three light stimulation scans lasting fifteen minutes each [Figure 1B]. After every scan, the participants were asked six questions about their subjective experiences (see below) and verbally responded via the speaker system of the scanner. An anatomical scan was performed before participants were released from the scanner and experiment.

### FLS-induced phenomenology

Phenomenological aspects of the FLS-induced state were retrospectively assessed using six questions of the Altered States of Consciousness Rating Scale (ASC-R; (Dittrich, 1998)), which were previously identified as most characteristic of the subjective experience (Bartossek et al., 2021). The questions were applied in German, taken from original version of the 5D-ASC (Dittrich, 1998) and participants were asked to rate by verbally naming a value from 0-100% for how much the following statements apply: (1) *Ich fühlte mich schläfrig* English: *I felt sleepy,* (2) *Ich fühlte mich körperlos* English: *I had the impression I was out of my body,* (3) *Wie im Traum waren Raum und Zeitgefühl verändert* English: *My sense of time and space was altered as if I was dreaming*, (4) *Ich fühlte mich wie in einer wunderbaren anderen Welt* English: *I felt I was in a wonderful other world* (5) *Ich sah regelmäßige Muster*, English: *I saw regular patterns* (Note: In the original version the statement continues as: *… with closed eyes or in complete darkness*) (6) *Ich sah Farben vor mir* English: *I saw colors* (Note: In the original version the statement continues as: *… with closed eyes or in complete darkness*). We ran one-way repeated-measures ANOVAs to test the effect of experimental condition on ASC-R questionnaire ratings using the *rstatix* package in Rstudio (v2022.07.2). As distribution of ratings had a tendency for skewedness (e.g., positive skew for pre and post scans), significant ANOVA results were additionally confirmed using non-parametric Kruskal-Wallis testing.

### fMRI Scanning

Participants were scanned using a 3T Siemens Tim Trio MRI scanner equipped with a 32-channel head coil (Siemens Medical, Erlangen, Germany). For resting-state fMRI images, a T2*-weighted echo planar imaging (EPI) sequence was used (37 axial slices acquired interleaved, in-plane resolution is 3mm^2^, slice thickness = 3 mm, flip angle (FA) = 70°, 20% gap between slices, repetition time (TR) = 2000 ms, echo time (TE) = 30 ms). A structural image was acquired for each participant using a T1-weighted image acquired with Magnetisation prepared rapid gradient-echo (MPRAGE) sequence (TR = 1900 ms, inversion time = 900 ms, TE = 2.52 ms, FA = 9°, voxel size 1mm^3^). Head motion was minimized using cushioned supports to restrict movement.

### MRI data pre-processing

Data were pre-processed and analysed using a custom-built resting-state data analysis pipeline within SPM12 (www.fil.ion.ucl.ac.uk/spm/). The anatomical T1-images were normalized to MNI152 space using the segmentation approach, by estimating a nonlinear transformation field, which is then applied to the functional images. Slice time correction and realignment was applied to the functional data before spatial normalisation to MNI152 space using unified segmentation in SPM12, which includes reslicing to an isometric 2 mm voxel size (Ashburner and Friston, 2005). The frame-wise displacement (FD) was calculated for each scan using BRAMILA tools (Power et al., 2012). Volumes that exceeded a threshold of 0.4 mm were masked during following analysis steps (*“scrubbing”*). Principal component analysis (CompCor) was done using the DPABI toolbox (toolbox for Data Processing & Analysis of Brain Imaging, http://rfmri.org/dpabi) within the CSF/white matter mask on the resting-state data to estimate nuisance signals (Behzadi et al., 2007). Anatomical masks for CSF, white and grey matter were derived from tissue-probability maps provided in SPM12. Smoothing was performed with a 3 mm FWHM Gaussian kernel, to retain high spatial specificity of small ROIs within the thalamus. The first five principal components of the CompCor analysis, six head motion parameters, linear and quadratic trends as well as the global signal were used as nuisance signals to regress out associated variance. The removal of global signal changes has been controversially discussed in resting-state fMRI literature with arguments for and against (see (Murphy and Fox, 2017) for overview). It has been particularly discussed for pharmacological studies (e.g., (Carhart-Harris et al., 2012; Vollenweider and Preller, 2020)), in which changes in blood flow, blood pressure, breathing rate and other physiologic parameters might account for some aspects of ROI-to-ROI correlations. Until conclusive interpretations of such differences are revealed, it is suggested to report data with and without global signal regression (Vollenweider and Preller, 2020). Therefore, we additionally report all analyses without GSR in the Supplement. Finally, the toolbox REST (www.restfmri.net) was used for temporal band-pass filtering (0.01-0.08 Hz).

### ROI-to-ROI correlation analysis

We used the AAL3 (Rolls et al., 2020) to define anatomical ROIs for thalamus parcellation (14 thalamic nuclei for each hemisphere) and a volume-based MPM of visual topography for cortical parcellation (23 visual areas for each hemisphere; see Figure 1C) (Wang et al., 2015). Using probability maps of V1 and V2, overlapping ROIs at the midline were resolved by assigning voxels to the region with highest probability. Of thalamic ROIs, the Reuniens nucleus is only 8mm^3^ and was not included in our analyses.

For each ROI, mean BOLD time courses were extracted and temporal ROI-to-ROI correlations calculated. For all ROI-to-ROI pairs, we averaged the correlation coefficients of pre and post scans and then computed differences with experimental conditions via subtraction of matrices. We took the mean of correlation coefficients for ipsilateral connections (e.g., left LGN and left V1v averaged with right LGN and right V1v) to give one bilateral functional correlation coefficient for each pair of ROIs. Lilliefors test of normality showed that at least 90% of ROI-to-ROI correlation coefficients were Gaussian distributed across participants for each condition. Therefore, to test for specific changes within thalamus and visual areas, we ran repeated-measures ANOVAs with condition as a fixed effect and connectivity changes as the dependent variable. We selected 16 visual areas to test: 8 within the ventral stream (V1v, V2v, V3v, hV4, VO1, VO2, PHC1, PHC2) and 8 within the dorsal stream (V1d, V2d, V3d, V3a, V3b, LO1, LO2, hMT), as classified by Wang *et al*. (2015). We conducted the analyses for LGN, inferior and lateral pulvinar. We Bonferroni-corrected the alpha threshold to .003 (.05/16) to correct for 16 repeated-measures ANOVAs for every thalamic nucleus. When tests were significant, post-hoc t-tests were used to determine the differences between condition groups. Thereafter, we further explored the ROI-to-ROI connectivity matrices of all thalamic and visual ROIs to identify if functional connectivity with any other thalamic nuclei or visual areas appeared to be modulated by FLS.

### Testing the relationship between subjective experience and connectivity changes

To test for the relationship between subjective experience and connectivity changes, we selected ratings of “I saw regular patterns” and “I saw colours” to reflect the intensity of visual phenomena. Using paired t-tests, we tested whether the distribution of ratings between the two items were different for each condition. As these tests were nonsignificant, we took the average of seeing patterns and seeing colours for each participant as the measure for occurrence of visual hallucinations. We subtracted the average of pre and post ratings from those of each experimental condition. From here, we ran linear mixed effects models with change in subjective ratings as a fixed effect and change in functional connectivity as the dependent variable. Participants were included as a random effect. Models with random intercept only had better fit (i.e., lower Akaike Information Criterion values) than random intercept and random slope models, and therefore models were run with random intercepts only. Following our hypotheses, we ran this test for connectivity changes between LGN, inferior pulvinar, lateral pulvinar and 16 visual subregions, thus Bonferroni-correcting the alpha threshold to .003. The analyses were conducted using the *lme4* package in Rstudio (v2022.07.2). Underlying assumptions of linear mixed modelling (e.g., equal variance of residuals, Gaussian-distributed dependent variable) were tested and met.

## Results

### FLS-induced subjective experience

We assessed the subjective experience following each scanning session (including pre/post scans) to test whether reported effects induced by the experimental conditions were comparable to previous findings where FLS was applied outside of the MRI (Amaya et al., 2023; Bartossek et al., 2021). Using ASC-R scores from each session, we ran 5x1 repeated-measures ANOVAs to test the effect of condition on ASC ratings. The results are presented in Figure 2. We found a significant effect of experimental condition on ratings of “I felt I was in a wonderful other world” (F(4, 92) = 9.83, *p* < .001), “My sense of time and space was altered as if I was dreaming” (F(4, 92) = 7.12, *p* < .001), and “I had the impression I was out of my body” (F(4, 92) = 5.37, *p* < .001), where post-hoc paired t-tests revealed that 10 Hz elicited significantly higher ratings than pre and post resting scans (*p*<.05). Further, there was a significant effect of experimental condition on ratings of “I saw patterns” (F(2.09, 48.03) = 104.53, *p* < .001) and “I saw colours” (F(2.2, 50.71) = 88.78, p < .001), where post-hoc paired t-tests showed that all experimental conditions were significantly different from each other and 10 Hz generated the highest ratings (*p* < .001) [Figure 2]. Non-parametric Kruskal-Wallis testing confirmed all ANOVA results (Out of body: H(4) = 14.23, *p* = .006; Altered time and space: H(4) = 15.88, *p* = .003; Wonderful other world: H(4) = 16.79, *p*= .002; Patterns: H(4) = 87.61, *p*<.001; Colours: H(4) = 87.84, *p* < .001), together showing that FLS inside the MRI scanner robustly induced hallucinatory experiences in all participants, with 10 Hz stimulation eliciting the highest intensity of subjective experience

**Figure 2.**
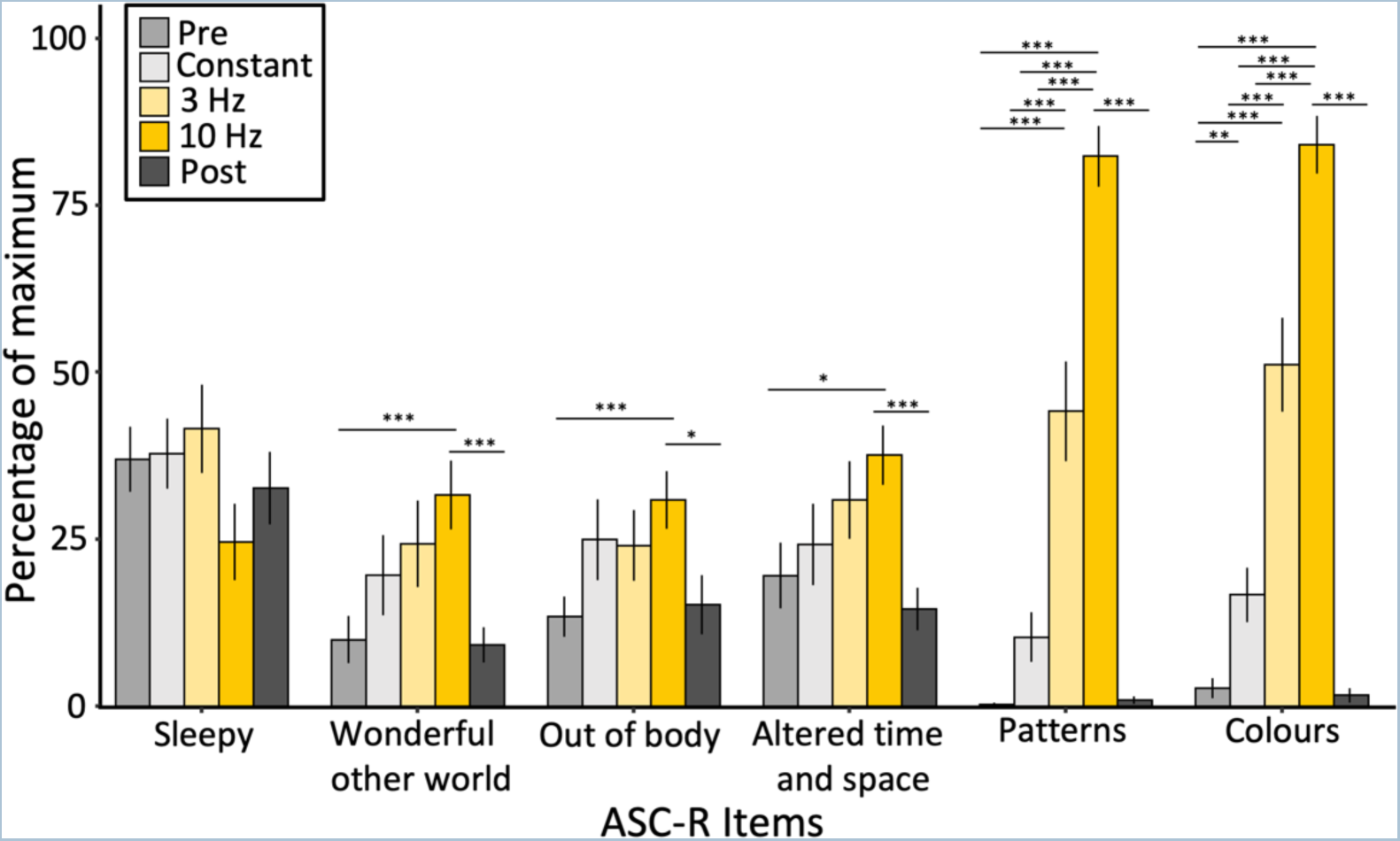
Mean scores of ASC-R items for each experimental condition, indicating no differences on wakefulness across conditions, minor parametric effects on general ASC phenomena and a strong modulation of visual phenomena. Effects tested via one-way repeated-measures ANOVAs; post-hoc paired t-test significance represented by * *p*<.05, ** *p*<.01, *** *p*<.001. Standard error is depicted by error bars.

### Changes in functional connectivity between LGN and visual areas

We tested for effects of FLS (3 Hz, 10 Hz) and constant light on connectivity changes from baseline between LGN nuclei and 16 visual areas using repeated-measures 3x1 ANOVAs (note that for every participant we averaged connectivity changes across ipsilateral connections; see methods). Alpha is Bonferroni corrected to .003 (.05/16). We found an increase in connectivity strength for 3 Hz and 10 Hz compared to baseline (average of pre and post scans), however 3 Hz and 10 Hz were not different from each other [Figure 3]. Specifically, there was a significant effect of experimental condition on connectivity changes between LGN and V1v (F(2, 46) = 17.75, *p* < .001), V1d, F(2, 46) = 20.64, *p* < .001), V2v (F(2, 46) = 26.91, *p* < .001), V2d (F(2,46) = 27.17, *p* < .001), V3v (F(2,46) = 20.73, *p* < .001), V3d (F(2,46) = 16.39, *p* < .001), which encompasses all early visual areas. In addition, there was a significant effect of experimental condition on connectivity changes with hV4 (F(2, 46) = 13.15, *p* < .001), VO1 (F(2, 46) = 7.40, *p* = .002) and V3a (F(2, 46) = 7.85, *p* = .001), whereby 10 Hz FLS induces the strongest coupling, followed by 3 Hz while constant light induced a weak decoupling.

**Figure 3.**
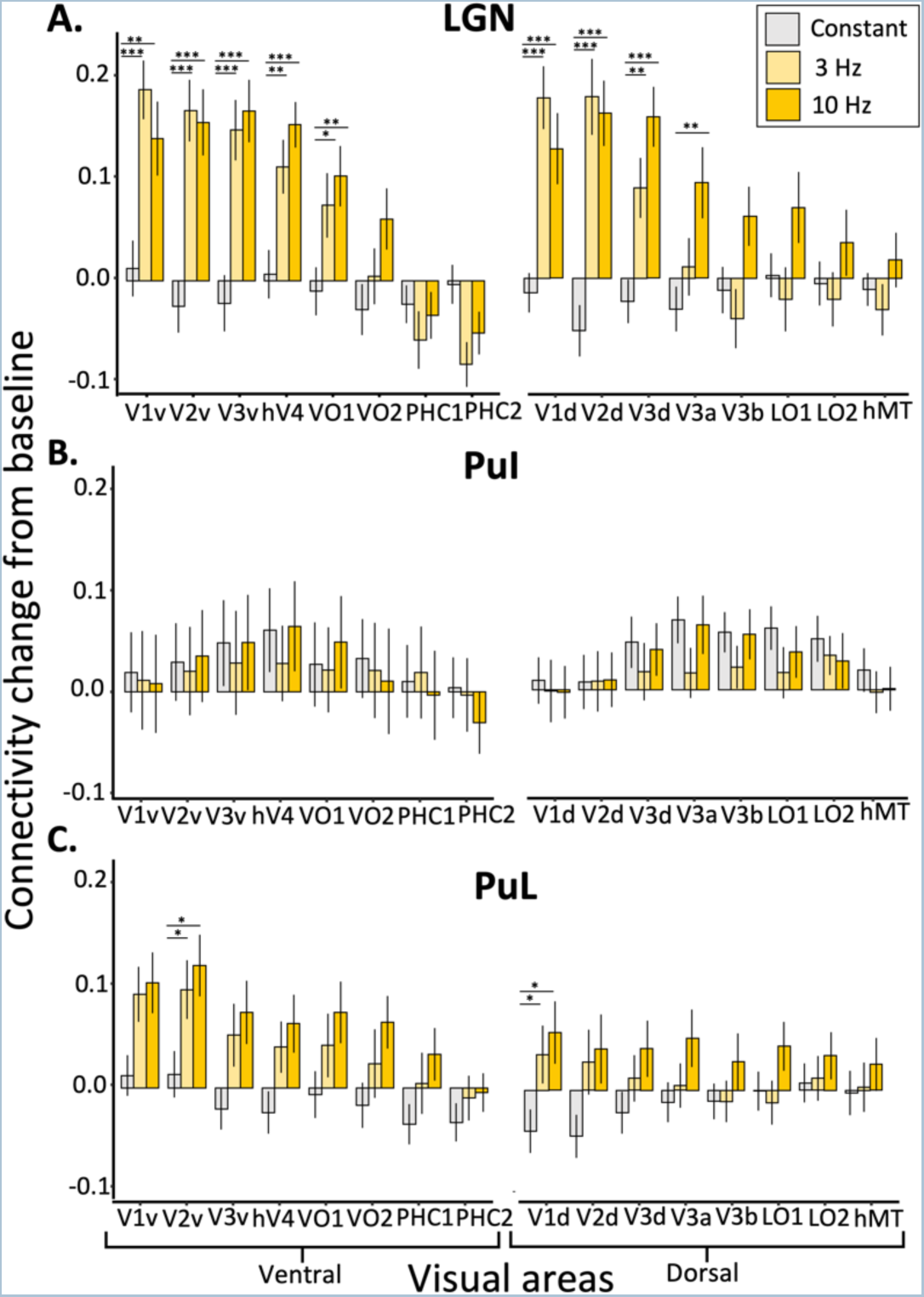
Effects of FLS on functional connectivity changes compared to baseline (average of pre and post scans) between visual areas and (A) LGN, (B) inferior pulvinar (PuI) and (C) lateral pulvinar (PuL). Visual areas are grouped into ventral and dorsal visual streams, as presented by Wang *et al*. (2015). Of the repeated-measures ANOVAs that returned significant effects of condition on connectivity change from baseline (alpha = .003), post-hoc paired t-tests indicate differences between conditions, where significance is represented by * *p*<.05, ** *p*<.01, *** *p*<.001. There is a strong modulation of condition on connectivity increases between LGN and EVC, and proximal upstream visual areas of dorsal (V3a) and ventral (hV4, VO1) streams. 3 Hz and 10 Hz induce LGN hyperconnectivity to the same degree for EVC, however for higher areas of the dorsal stream (V3a, V3b, LO1), LGN hyperconnectivity is only apparent during 10 Hz FLS. There is no significant effect of light stimulation on connectivity changes between inferior pulvinar and visual areas, while lateral pulvinar shows a similar pattern of connectivity changes as LGN with visual areas, albeit less strong.

### Changes in functional connectivity between pulvinar and visual areas

Using the same treatment of data as for the LGN, we tested for effects of experimental condition on connectivity changes between inferior and lateral divisions of the pulvinar and 16 subregions of the visual cortex. Using repeated-measures 3x1 ANOVAs, we found no significant effects of condition on connectivity changes with inferior pulvinar across all tested visual areas, which is evident in Figure 3. For the lateral pulvinar, a significant effect of condition was revealed for connectivity changes with V1d (F(2,46) = 7.13, *p* = .002) and V2v (F(2,46) = 7.47, *p* = .002), whereby post-hoc t-tests showed that 10 Hz and 3 Hz induced stronger coupling than constant light (*p* < .05) [Figure 3].

### Association of connectivity strength with subjective experience

We ran linear mixed models with rating as a fixed effect and participant-specific random intercepts to determine if ASC-R mean ratings of experienced visual effects (i.e., mean of “I saw patterns” and “I saw colours”; see Methods) could predict changes in functional connectivity between ROIs. Alpha was Bonferroni-corrected to .003 to account for the comparison of 16 models for each group (i.e., 16 visual areas for LGN, lateral pulvinar and inferior pulvinar). We found that subjective ratings significantly predicted increases in connectivity between the LGN and V2v (*p* < .001; R^2^m = 0.15; R^2^c = 0.46), V2d (*p* < .001; R^2^m = 0.15; R^2^c = 0.44), V3v (*p* < .001; R^2^m = 0.19, R^2^c = 0.54), V3d (*p* < .001; R^2^m = 0.26, R^2^c = 0.59), hV4 (*p* < .001; R^2^m = 0.22, R^2^c = 0.50), VO1 (*p* = .002; R^2^m = 0.12, R^2^c = 0.55), VO2 (*p* = .001; R^2^m = 0.10, R^2^c = 0.48), V3a (*p* < .001; R^2^m = 0.16, R^2^c = 0.58) and V3b (*p* = .001; R^2^m = 0.10, R^2^c = 0.46). Furthermore, subjective ratings significantly predicted connectivity increases between lateral pulvinar and V1d (*p* < .001; R^2^m = 0.10; R^2^c = 0.56). Together, this shows that the subjective ratings associate moreso with LGN interactions with upstream visual areas beyond V1, while connectivity between V1 and lateral pulvinar are significantly associated with subjective ratings.

### Exploratory analysis of thalamocortical connectivity

To explore the whole connectivity profiles of thalamic nuclei, we plotted connectivity matrices with 14 thalamic ROIs from the AAL3 atlas (Rolls et al., 2020) and 23 cortical ROIs from the Wang *et al*. maximum probability map of visual topography (Wang et al., 2015). Figure 4 displays the ROI-to-ROI correlation coefficients of the experimental conditions subtracted by the averaged pre and post scans, thus showing the connectivity change induced by the conditions. We see strong hyperconnectivity between AV, ventral and MD thalamic nuclei and cortical visual regions. Connectivity with higher-order cortical visual regions, such as hV4, VO1 and LO1, show more frequency-dependent effects (i.e., 10 Hz induces more coupling than 3 Hz) than EVC. We note that, as ventral nuclei (i.e., VA, VPL and VL nuclei) show similar changes in connectivity patterns and have anatomical proximity, we consider their effects collectively as a ventral group. Likewise, MDM and MDL are divisions of MD nuclei showing similar connectivity patterns and are therefore considered together as the MD region. The changes in connectivity induced by 10 Hz FLS are additionally represented in Figure 6.

**Figure 4.**
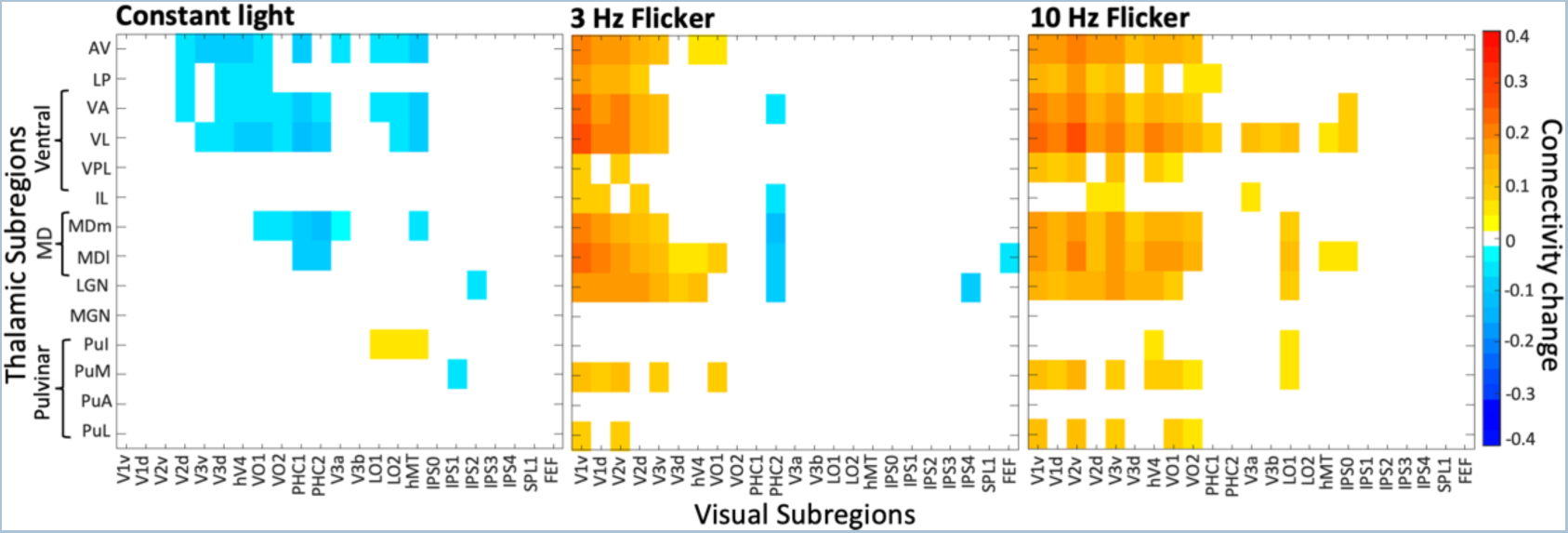
Connectivity matrices for all visual areas and thalamic nuclei during constant light, 3 Hz and 10 Hz FLS, subtracted by the average of pre and post scans (closed-eye rest) to represent the connectivity change induced by the experimental conditions. A mask has been applied where only significant connectivity changes compared to baseline are shown, as determined by paired t-tests (alpha=.01). All divisions of ventral nuclei form a cluster as they display similar connectivity patterns. Likewise, medial and lateral divisions of MD nuclei show similar connectivity patterns and can be collectively considered as the MD region. With this clustering, we observe that AV, ventral and MD thalamic regions display the greatest frequency-dependent effects of FLS on connectivity changes with visual areas, in that 10 Hz induces the strongest coupling that is additionally evident in upstream visual areas of both ventral (e.g., VO2) and dorsal (e.g., hMT, V3a) visual streams. Meanwhile, we see overall hypoconnectivity in the constant light condition.

**Figure 6.**
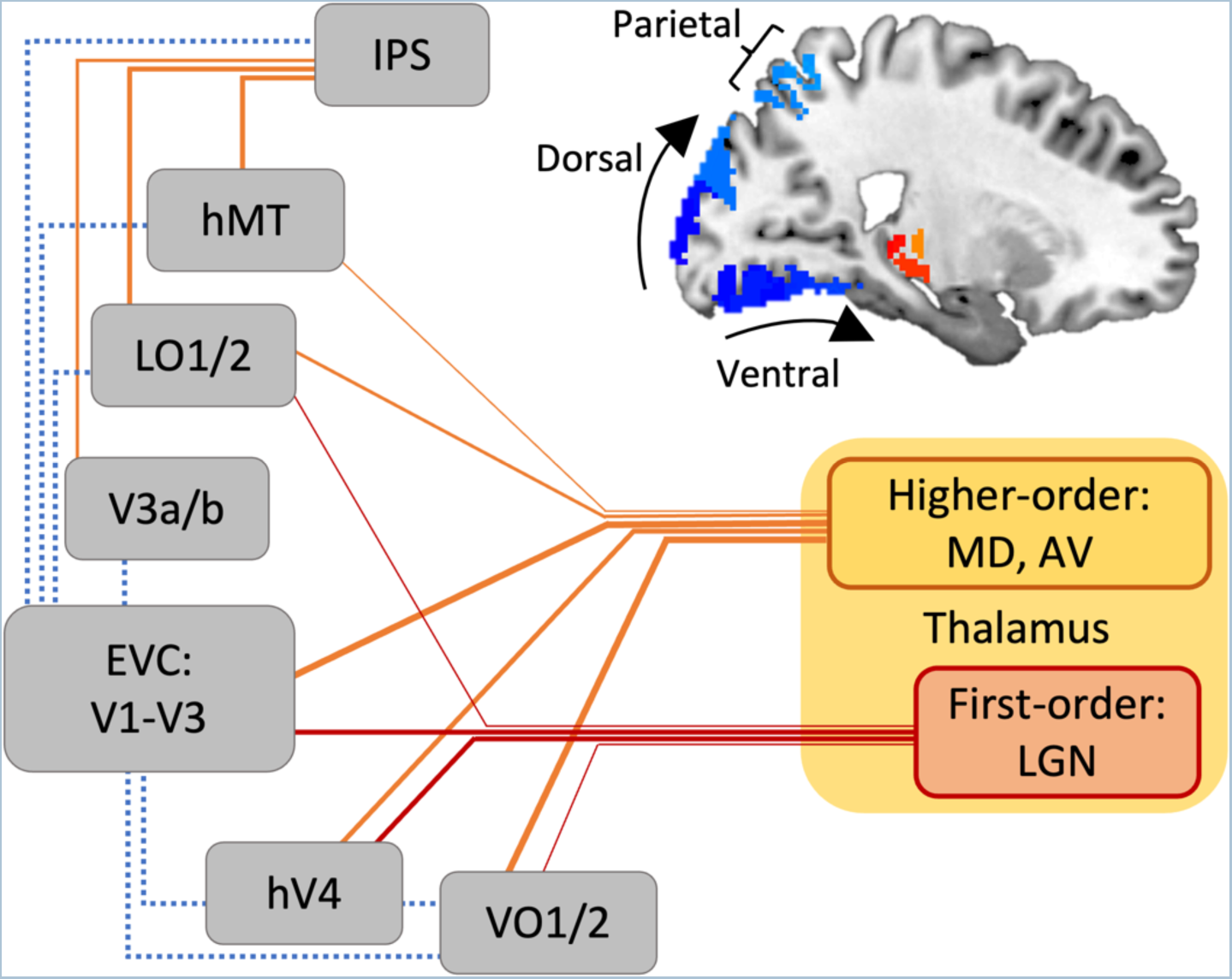
Summary of thalamocortical and corticocortical functional connections changes during 10 Hz FLS. Orange lines represent functional hyperconnectivity compared to baseline, while blue dashed lines represent hypoconnectivity. For display purposes, red and orange are used to distinguish between first-order (sensory) and higher-order (non-sensory) thalamic regions. The strength of correlation change is depicted by line thickness. During FLS, the LGN shows connectivity increases to early visual cortices (EVC; V1-V3) and proximal upstream areas of the ventral stream (i.e., hV4, VO1). We found that AV and MD higher-order thalamic nuclei display increased coupling with visual areas along ventral and dorsal visual streams. (Note: ventral nuclei also displayed a comparable connectivity profile, while their contributions as higher-order nucleus are less clear). Connectivity changes of higher-order nuclei with ventral areas are notably stronger than LGN coupling. As these nuclei do not receive direct driving retinal inputs, they are most likely driven by inputs from EVC. While the directionality of effects in higher-order regions are speculative, our findings may indicate that AV and MD nuclei take an orchestrating role for information flow across cortical regions, such as eliciting the observed hypoconnectivity between EVC and upstream cortical areas.

### Exploratory analysis of visual area and thalamic interconnectivity

To explore interconnectivity profiles of visual and thalamic areas, we plotted masked interconnectivity matrices of 23 cortical and 14 thalamic visual ROIs, where only connectivity changes that were significantly different from baseline (alpha = .01) are displayed [Figure 5]. Figure 5A shows that 3 Hz and 10 Hz FLS leads to hyperconnectivity within EVC (i.e., V1-V2) but hypoconnectivity between EVC and higher visual areas of both the ventral (e.g., LO2/1) and dorsal (e.g., V3a/b, IPS) streams. Meanwhile, higher visual areas show increased coupling to each other (e.g., LO2/1 and IPS). Interconnectivity matrices of thalamic ROIs display few changes in connectivity amongst thalamic nuclei, especially at 10 Hz FLS [Figure 5B].

**Figure 5.**
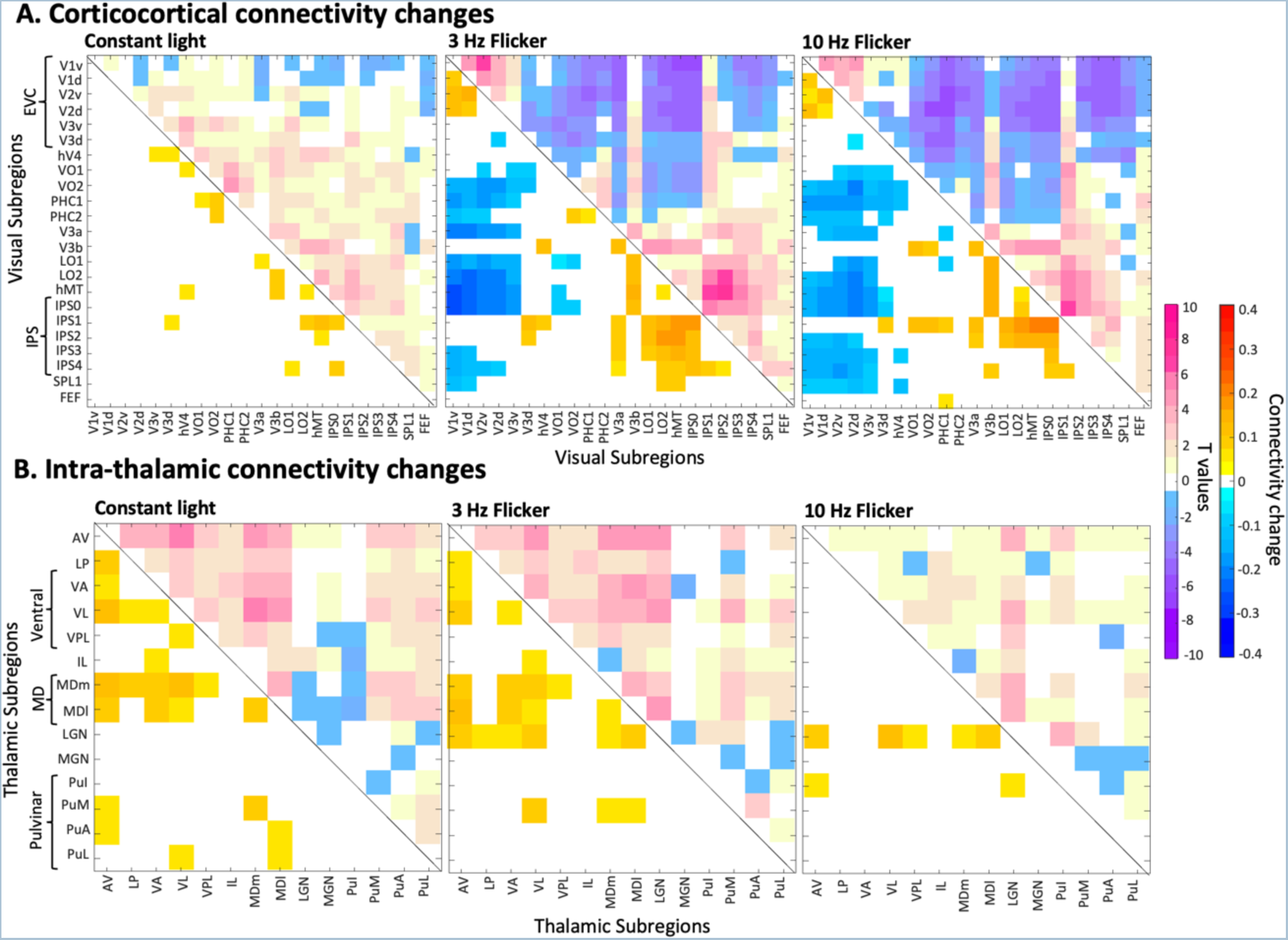
(A) Connectivity changes between visual areas during constant light, 3 Hz and 10 Hz FLS, as compared pre and post scans. Upper half of matrices show t values of paired t-tests; lower half show connectivity changes (masked at *p*<.01, where paired t-tests revealed a significant difference between connectivity in experimental condition versus baseline). There are two groups of hyperconnectivity: within EVC and between upstream visual areas (e.g., LO1, hMT) and IPS, while these groups are decoupled from each other. (B) Connectivity changes within the thalamus. The connectivity changes in the 10 Hz condition are confined to relevant areas (i.e., LGN, AV, ventral and MD nuclei), which supports that the applied parcellation yields region-specific effects. If ROIs were to reflect the same underlying signals, one would expect an overall increase in connectivity between thalamic subfields. Thalamo-cortico-thalamic connections likely drive increased LGN connectivity with other thalamic nuclei (i.e., AV, ventral and MD nuclei), such that the signal passes from LGN via visual cortices to higher-order thalamic nuclei.

## Discussion

We tested the effects of FLS on functional connectivity between anatomically specified thalamic nuclei and visual areas. We found that FLS induced hyperconnectivity between the LGN and early visual cortices (EVC: V1-V3), independent of flicker frequency. Meanwhile, upstream visual areas show a differential effect of flicker frequency on LGN connectivity, in that coupling was strongest for 10 Hz. Similarly, FLS induced a frequency-dependent increase in participant ratings of visual hallucinations (“I saw colours” and “I saw patterns”), which replicates previous findings (Amaya et al., 2023; Bartossek et al., 2021). The intensity of visual phenomena was associated with the strength of connectivity changes between LGN and higher visual areas, especially for V3 and hV4, suggesting that effects are not only driven by a simple feedforward mechanism from LGN to V1, but rather arise from a modulation of upstream visual areas. FLS additionally induced weak thalamocortical hyperconnectivity with the lateral pulvinar but had no effect on the inferior pulvinar. Hyperconnectivity between lateral pulvinar and V1 was associated with subjective ratings, which may correspond to a top-down modulatory influence of the pulvinar on V1. When exploring connectivity changes across all thalamic nuclei of the AAL3 atlas, we found stronger frequency-dependent modulations of connectivity between AV, ventral and MD thalamic nuclei and visual areas. Moreover, we explored corticocortical connectivity changes between visual areas and observed two groups of hyperconnectivity: (1) within EVC and (2) between upstream visual areas and intraparietal sulcus (IPS), while these groups were decoupled from each other. Overall, we identify that hyperconnectivity between upstream visual areas, LGN and other thalamic regions, such as AV, ventral and MD nuclei, may be most relevant for the emergence of visual hallucinations.

### Thalamocortical connectivity with LGN

FLS significantly increased connectivity between LGN and EVC, hV4, VO1 and V3a. This expected finding supports that rhythmic retinal activation propagate along dorsal (i.e., V3a) and ventral (i.e., hV4, VO1) visual streams. It is likely that driving inputs from the retina cause synchronisation with LGN and subsequent visual areas via excitatory feedforward signalling, which manifests as an increase in functional connectivity, in the sense of entrainment. EEG studies have shown that periodic visual flicker at alpha frequency increases neural entrainment at that frequency (Adrian and Matthews, 1934; Mathewson et al., 2012; Notbohm et al., 2016; Notbohm and Herrmann, 2016; Schwartzman et al., 2019), which coincides with findings that subjectively experienced FLS-effects are strongest in the alpha-frequency range (Amaya et al., 2023).

As there are rich feedback connections between visual cortices and thalamic nuclei (Budd, 2004; Murphy et al., 1999), an increase in functional connectivity likely encapsulates both feedforward and feedback processes. Indeed, it was recently shown that visual flicker induced phase-locking in LGN and cortical layers 4 and 5 of V1 (Schneider et al., 2023), which are involved in feedforward and feedback processes, respectively. The corticogeniculate inputs may refine the feedforward signals, possibly through enhancing response precision and synchronising LGN action potentials (Andolina et al., 2007; Briggs, 2020; Sillito et al., 1994), leading to the development of specific geometric patterns and distinct colours. While our data may represent changes to both feedforward and feedback interactions during FLS, future research should assess the weighting of these contributions to the resulting thalamocortical hyperconnectivity.

### Thalamocortical connectivity with pulvinar

Due to the involvement of the pulvinar as a higher-order thalamic nucleus in visual processing (Adams et al., 2000; Benevento and Rezak, 1976; Guedj and Vuilleumier, 2020; Kaas and Lyon, 2007), we expected to find effects of FLS on thalamocortical connectivity with the inferior and lateral pulvinar. The lateral pulvinar demonstrated increased coupling with EVC, which was more apparent for ventral visual areas compared to dorsal (see Figure 3). This reflects the major contribution of the lateral pulvinar to the ventral visual stream (Kaas and Lyon, 2007), which is responsible for shape and colour recognition (Ungerleider and Haxby, 1994), possibly relating to the hallucinatory perception of patterns and colours, although tests of this association did not survive conservative Bonferroni correction. Additionally, there were no effects of FLS on inferior pulvinar connectivity, together showing that the effects of FLS on pulvinar connectivity were smaller than expected, especially when compared to other thalamic nuclei (see below). It is possible that the observed effects on the pulvinar can be assigned to contributions to visual attention (Gattass et al., 2017; Saalmann et al., 2012), rather than the subjective experience of visual hallucinatory phenomena.

### Further thalamic nuclei displaying altered thalamocortical connectivity

Exploratory analyses of ROI-to-ROI connectivity highlighted three further thalamic subregions whose connectivity to visual areas seem to be modulated by FLS: (1) anterior nuclei, where all divisions of anterior nuclei are included in the AV region of the AAL3 atlas. (2) the ventral nuclei group, which includes VA, VPL and VL nuclei, and (3) MD nuclei. All these thalamic regions showed hyperconnectivity with EVC during 3 Hz and 10 Hz FLS and hyperconnectivity with further upstream visual areas (e.g., V3a, VO2) for 10 Hz FLS only.

Anterior nuclei are criycally involved in spayal navigayon and memory (Roy et al., 2022; Safari et al., 2020). For example, they receive head direcyon signals through vesybular sensory inputs (Peyrache et al., 2019; Sharp et al., 2001; Taube, 2007). Within the anterior division, AV nuclei are linked to the visual cortex via connecyons to the retrospinal cortex (Lomi et al., 2023), which is thought to contribute to spayal organizayon in imaginayon (Botzung et al., 2008; D’Argembeau et al., 2008; Hassabis et al., 2007; Szpunar et al., 2007). Furthermore, a post-mortem study of payents with schizophrenia found fewer thalamocorycal projecyons in the AV nucleus bilaterally (Danos et al., 1998), suggesyng a potenyal role in pathologic altered perceptual processing.

Within the ventral group, the VL nucleus has been associated with auditory-tactile synaesthesia (Ro et al., 2007), despite being primarily known as a first-order relay for motor inputs (Percheron et al., 1996). The ventral thalamic group were found to be hyperconnected with sensorimotor networks within psychosis (Avram et al., 2018) and following LSD administration (Avram et al., 2022), together indicating an contribution to altered perceptual processing. VL nuclei were further found to be functionally connected to the lateral visual network (Kumar et al., 2022), which is involved in motion and shape perception (Smith et al., 2009). Here, despite being known as first-order nuclei, we speculate that ventral thalamic regions may serve higher-order, integrative functions within visual processing, as it was found that first-order nuclei can also form a hub for interactions with multiple functional networks (Hwang et al., 2017).

MD nuclei have extensive connections with the prefrontal cortex (Haber and Mcfarland, 2001) and are primarily involved in executive cognitive function (Parnaudeau et al., 2017). For patients with psychotic disorders, MD nuclei were functionally hypoconnected with prefrontal areas (Avram et al., 2018; Woodward and Heckers, 2016) while being hyperconnected with sensorimotor areas (Anticevic et al., 2014). Further, in a healthy population, MD nuclei were activated during perception of fused versus non-fused colour (indicative of hallucinatory perception; (Seo et al., 2022)). Ventral and MD nuclei were also highlighted in recent reviews of relevant thalamic regions contributing to drug- and pathology-related hallucinatory phenomena (Avram et al., 2021; Doss et al., 2021).

Together, these thalamic regions (AV, ventral, MD) show a similar thalamocortical hyperconnectivity pattern with visual cortices as recently reported in patients with chronic schizophrenia (Rolls et al., 2021), suggesting that the observed hyperconnectivity could be associated with hallucinatory experiences. However, the sparsity of literature linking these thalamic regions to the visual system makes it difficult to infer how exactly they contribute mechanistically to the emergence of visual hallucinations. This calls for further research into the functional involvement of higher-order nuclei in visual processing and consequently in hallucinatory experiences.

The role of higher-order thalamic nuclei in the formayon of visual hallucinayons may instead lie in their ability to orchestrate brain-wide corycal acyvity. AV, ventral and MD nuclei all display strong connector hub properyes for corycal funcyonal networks (Hwang et al., 2017), which suggests that these thalamic areas are not only funcyonally specific, but also contribute to domain-general, brain-wide funcyon (Shine et al., 2023). For example, cross-frequency coupling (CFC) may be a mechanism underlying FLS-induced effects, whereby the thalamus and/or EVC are entrained to alpha frequency, which consequently modulates large-scale corycal excitability occurring in the gamma frequency range (Canolty and Knight, 2010; Klimesch et al., 2007; Kosciessa et al., 2021; Wang et al., 2012). Given the importance of the thalamus in coordinating brain-wide activity, thalamocortical pathways have become a central feature of multiple theories of consciousness (Alkire et al., 2008; Aru et al., 2019; Purpura and Schiff, 1997; Tononi and Edelman, 1998; Ward, 2011) e.g., Dynamic Core Theory ((Tononi and Edelman, 1998); from which Integrated Information Theory developed (Tononi, 2011; Tononi et al., 2016)), where subjective experiences might partly depend on orchestrating roles of thalamocortical interactions. Further, Dendritic Integration Theory proposes that cortical layer 5p neurons, where dendritic signalling is under control of thalamocortical projections of higher-order nuclei, are critical for conscious experience (Aru et al., 2019). While our study does not directly address the neural mechanisms of conscious processing, it adds to an understanding regarding the role of thalamocortical interactions within visual experiences in the context of hallucinatory perception. Future research should continue to integrate how thalamocortical interactions contribute to conscious awareness and phenomenal characteristics of subjective experience.

### Connectivity within visual areas

When exploring connectivity changes within the visual system, we found a consistent pattern of connectivity changes for 3 Hz and 10 Hz FLS. There was hyperconnectivity within two groups: (1) within EVC and (2) between upstream visual areas (e.g., LO1) and the IPS, while these two groups were decoupled from each other. Such altered corticocortical connectivity may have been mediated by the thalamus, which can sustain and modulate corticocortical functional connectivity (Schmitt et al., 2017). This connectivity pattern contrasts with effects on the visual system elicited by ASC pharmacological interventions, i.e., LSD, where a hypoconnectivity within EVC and within lateral visual regions (e.g., hV4/hMT) was found (Bedford et al., 2023; Carhart-Harris et al., 2016; Müller et al., 2018), however methodological differences in defining visual areas make it difficult to draw direct comparisons. Within our study, FLS-induced EVC hyperconnectivity likely reflects the visual sensory inputs that drive the consequent hallucinatory effects, while hallucinatory effects of LSD are mediated by serotonergic agonism (Aghajanian and Marek, 1999). It can be further speculated that EVC hyperconnectivity may evoke decoupling between EVC and upstream visual areas, which then become hyperconnected to further upstream areas (i.e., IPS) as a compensatory response. However, what the functional relevance of this would be is largely unclear, especially as similar corticocortical connectivity patterns for 3 Hz and 10 Hz FLS suggest that the altered connectivity does not correspond directly to the visual experience. Further research should explore in detail whether there is a phenomenal correlate of altered connectivity along the visual hierarchy and furthermore, whether a temporal sequence of connectivity changes can hint towards a causational link.

### Limitations

It must be acknowledged that not all thalamic nuclei are accounted for in the AAL3 atlas. Particularly, the thalamic reticular nucleus, which forms a thin sheet surrounding the thalamus (Pinault, 2004) is known to exert inhibitory control on other thalamic nuclei, such as the LGN (Halassa and Sherman, 2019). Therefore, it is possible that changes in connectivity with thalamic regions may have been mediated by thalamic reticular activity. While future work could utilise other atlas parcellations of the thalamus that include reticular nuclei (e.g., thalamic probabilistic atlas (Iglesias et al., 2018)), Rolls *et al*. (2020) purposely omitted this ROI from their atlas due to its difficult structure for automated parcellation. Accurate parcellation of the reticular nuclei may only be possible with a higher field MRI scanner (i.e., 7 Tesla) and thus higher resolution images.

Moreover, with the quantification of functional connectivity, interpreting the directionality of effects is highly limited. The well-described anatomy of retinal inputs to the thalamus allows to draw some inference on feedforward signalling from the LGN to the cortex. Furthermore, the lack of direct retinal inputs to higher-order nuclei suggests that these are most likely driven by corticothalamic signalling. However, an interpretation of directionality beyond these are speculative. Future investigations could employ effective connectivity analyses, such as regression Dynamic Causal Modelling (rDCM) (Frässle et al., 2021, 2017), as used in a recent LSD study (Bedford et al., 2023), which allows to test for directionality based on predefined network models of interacting regions. Thus, analyses such as rDCM could give more mechanistic insights into the sources of connectivity changes across thalamocortical and corticocortical loops. This may, in turn, shed further light on the functional roles of relay and higher-order thalamic nuclei in the generation of visual hallucinatory phenomena.

### Conclusions

Overall, we show that FLS induces thalamocortical hyperconnectivity between LGN, EVC and proximal upstream areas of ventral and dorsal visual streams (i.e., hV4, VO1, V3a). Additionally, while only weak effects were found for the pulvinar, hyperconnectivity between other thalamic nuclei and visual areas were more apparent, i.e., mediodorsal, anterior and ventral nuclei. The hyperconnectivity between higher-order thalamic nuclei and upstream visual areas was only evident for 10 Hz FLS, which follows the parametric modulation of flicker frequency on subjective ratings of seeing patterns and colours. This suggests that, although thalamocortical hyperconnectivity with LGN may initially drive the FLS-induced effects, the subsequent cortical interactions with higher-order thalamic nuclei may be more relevant for the emergence of visual hallucinations. In sum, we identify, for the first time, the specific thalamic nuclei and visual areas that display altered connectivity during flicker-induced hallucinatory phenomena.

## Acknowledgements

We thank Light Attendance GmbH (Innsbruck, Austria) for generously providing a Lucia N°03 system free of charge.

## Data and code availability

All data will be shared upon contact to I.A. Code for MRI preprocessing and generating connectivity matrices will be made available at: https://github.com/ioannaamaya/FLS-rsfMRI.git. Any further information required to re-analyse the data presented in this article is available without restrictions upon request to I.A.

## Author contributions

**Ioanna A. Amaya**: Formal analysis, Data curation, Methodology, Visualization, Writing – original draft. **Marianna E. Schmidt**: Data curation, Software. **Marie T. Bartossek**: Data curation. **Johanna Kemmerer**: Investigation, Data curation, Methodology. **Evgeniya Kirilina**: Methodology, Resources. **Till Nierhaus**: Methodology, Formal analysis, Software, Supervision, Writing – review & editing. **Timo T. Schmidt:** Conceptualisation, Investigation, Project administration, Resources, Methodology, Software, Supervision, Visualisation, Writing – original draft, Writing – review & editing.

