## Supplementary Figures for "Flicker light stimulation induces thalamocortical hyperconnectivity with LGN and higher-order thalamic nuclei"

### Supplementary Information

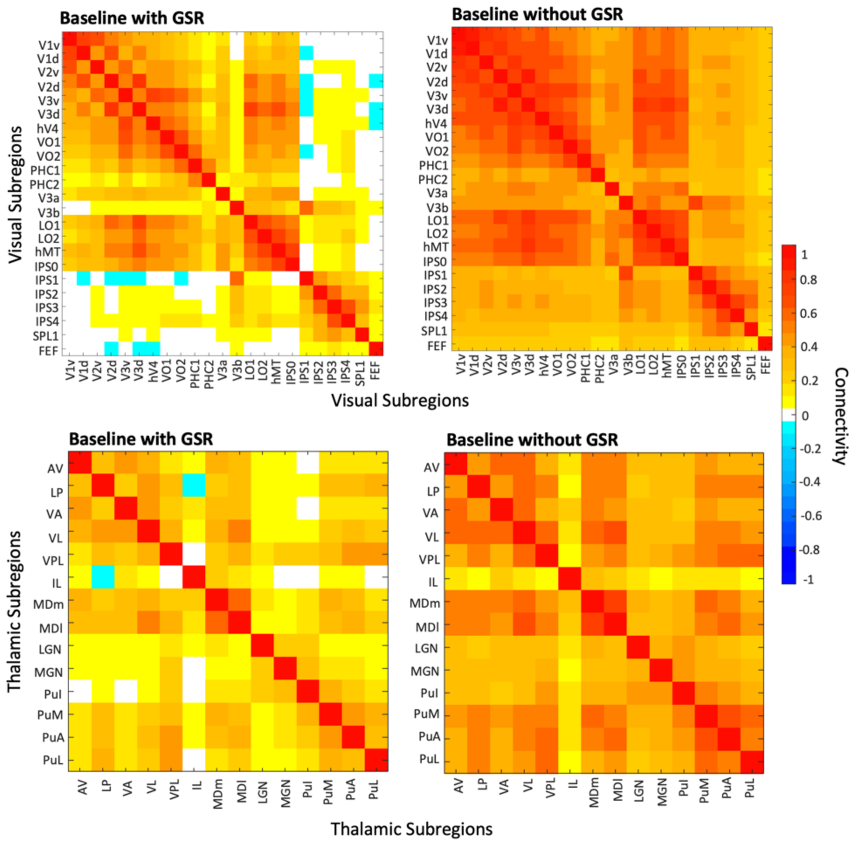


*Figure 1.* Intra-connectivity of (A) visual areas and (B) thalamic subregions with and without Global Signal Regression (GSR) during baseline, which constitutes the average of ROI-to-ROI correlational coefficients during pre- and post- resting-state closed eye scans.

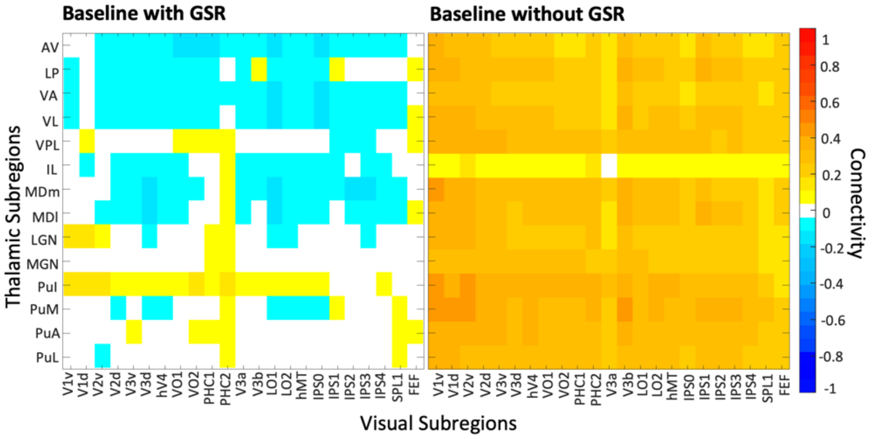


*Figure 2.* Thalamocortical connectivity during baseline (average of ROI-to-ROI correlational coefficients during pre- and post- resting-state closed eye scans) calculated with and without GSR.

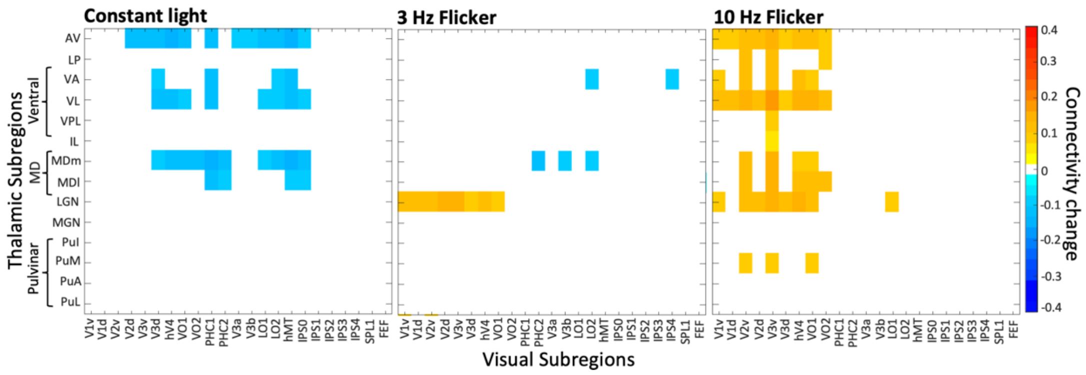


*Figure 3.* Connectivity matrices to show changes in thalamocortical connectivity without GSR for all visual areas and thalamic nuclei during constant light, 3 Hz and 10 Hz FLS, calculated by subtracting the average of pre- and post- scans from connectivity matrices during the experimental conditions. A mask has been applied whereby only significant connectivity changes compared to baseline are shown, as determined by paired t-tests (alpha=0.01).

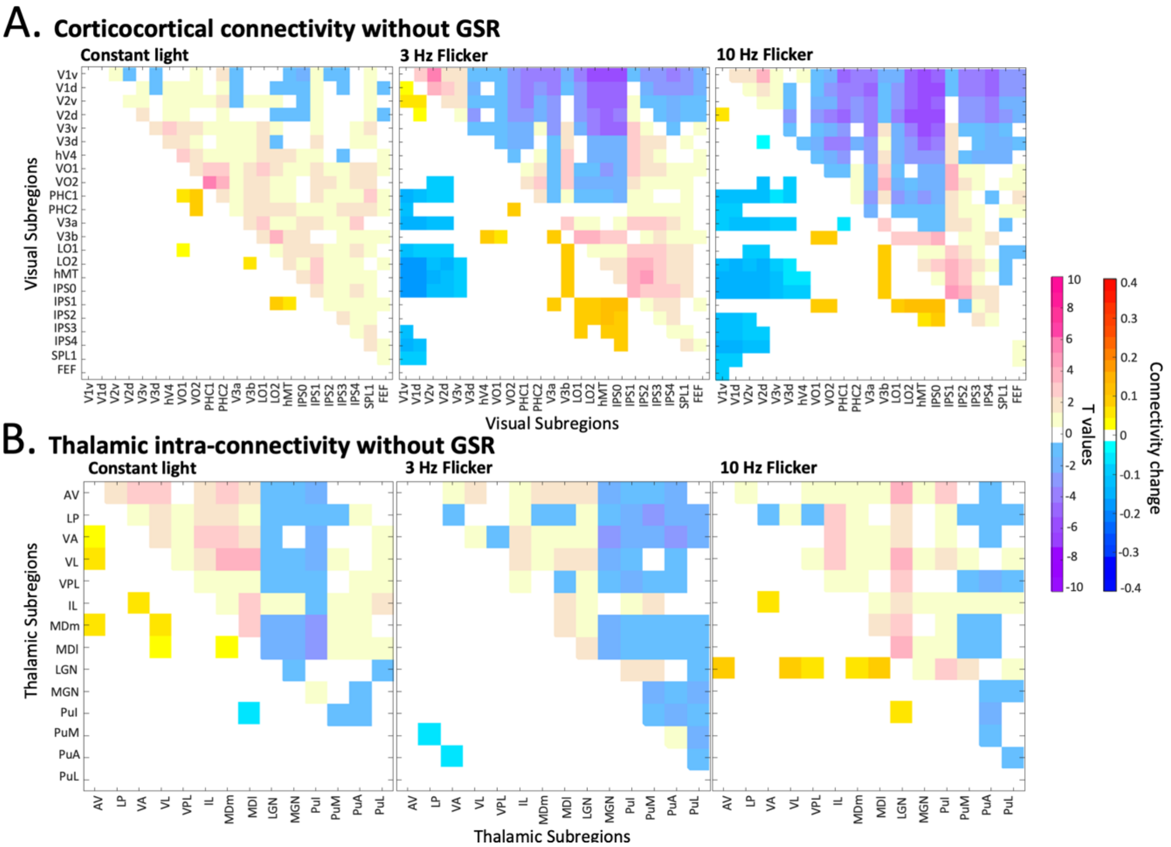


*Figure 4.* (A) Connectivity changes without GSR between visual areas during constant light, 3 Hz and 10 Hz FLS, as compared pre- and post- scans. Upper half of matrices show significance values of paired t-tests; Lower half show connectivity changes (masked at p<0.01). (B) Connectivity changes without GSR within the thalamus.
